# GxEMM: Extending linear mixed models to general gene-environment interactions

**DOI:** 10.1101/397638

**Authors:** Andy Dahl, Na Cai, Jonathan Flint, Noah Zaitlen

## Abstract

Gene-environment interaction (GxE) is a well-known source of non-additive inheritance. GxE can be important in applications ranging from basic functional genomics to precision medical treatment. Further, GxE effects elude inherently-linear LMMs and may explain missing heritability. We propose a simple, unifying mixed model for polygenic interactions (GxEMM) to capture the aggregate effect of small GxE effects spread across the genome. GxEMM extends existing LMMs for GxE in two important ways. First, it extends to arbitrary environmental variables, not just categorical groups. Second, GxEMM can estimate and test for environment-specific heritability. In simulations where the assumptions of existing methods do not hold, we show that GxEMM improves estimates of ordinary and GxE heritability and increases power to test for polygenic GxE. We then use GxEMM to prove that the heritability of major depression (MD) is reduced by stress, which we previously conjectured but could not prove with prior methods, and that a tail of polygenic GxE effects remains unexplained by MD GWAS.

## 1 Introduction

Gene-environment interaction (GxE) has been thoroughly documented at the level of individual genetic variants. In functional genomics, some variants have effects on expression that depend on external context [1–6], age [7], tissue [8], cell type [9, 10], or other genetic variants [11, 12]. In complex traits, GxE has been found for exposures like air pollution [13], child abuse [14], microbe exposure [15] and major lifetime stress [16]. Genetics can also interact with medical interventions, sometimes rendering treatment ineffective [17–19]. G-by-sex, perhaps the most studied form of genetic interaction, has been found for diseases including asthma, autism and diabetes [20–23].

GxE effects are among many explanations given for ‘missing heritability’, i.e. the gap between the variance explained by GWAS-significant SNPs and twin-based heritability estimates 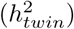 [24]. Although linear mixed models (LMMs) have shown that much of this missing heritability can be explained by many variants of small effect [25, 26], ordinary LMM heritability 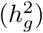 can still be substantially lower than 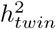. For example, major depression (MD) has 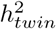 of roughly 30-40%, but 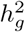 is more like 20-30% [27]. Because the linearity assumption inherent in LMMs excludes heritability derived from genetic interactions, GxE may explain some of this gap.

Despite these two parallel streams of work–seeking individual GxE effects and explaining missing heritability–comparatively little has been done in the intersection, i.e. to study GxE effects in aggregate across the genome. However, the success of ordinary LMMs suggests that GxE may be better understood at the polygenic level, by modeling genome-wide random effects, rather than by studying a few large-effect variants. Further, whether known GxE loci represent exceptions or the rule is under active debate for many complex traits, including MD, making genome-wide GxE inference valuable for basic research, experimental design, and precision treatment.

Several methods to estimate GxE genome-wide have already been developed. A useful heuristic, lacking a generative model, was proposed for the special case where environments are categorical and have equal heritabilities [28]. More recently, this has been generalized in several directions: to allow genetic and residual correlations across two contexts measured on all samples [29]; or to use only summary statistics [30]; or, for a single SNP, to allow high-dimensional environments [31].

We present a new, general form for polygenic gene-environment interactions that we call a GxE mixed model (GxEMM). While complex in some settings, GxEMM has a simple visual interpretation in the case of discrete environments (Figure 1). GxEMM generalizes and unifies several existing GxE mixed models, which provides theoretical value. But GxEMM’s primary value is pragmatic, as it extends the reach of GxE mixed models in two ways. First, it allows arbitrary environmental covariates, regardless whether they are continuous, discrete, proportions, or some combination. Second, GxEMM allows different environments to explain different amounts of GxE heritability; in particular, in the case of discrete environments, GxEMM can estimate and test for differential environment-specific heritability. Computationally, GxEMM uses existing software and thus has the same cost as generic REML methods.

**Figure 1:**
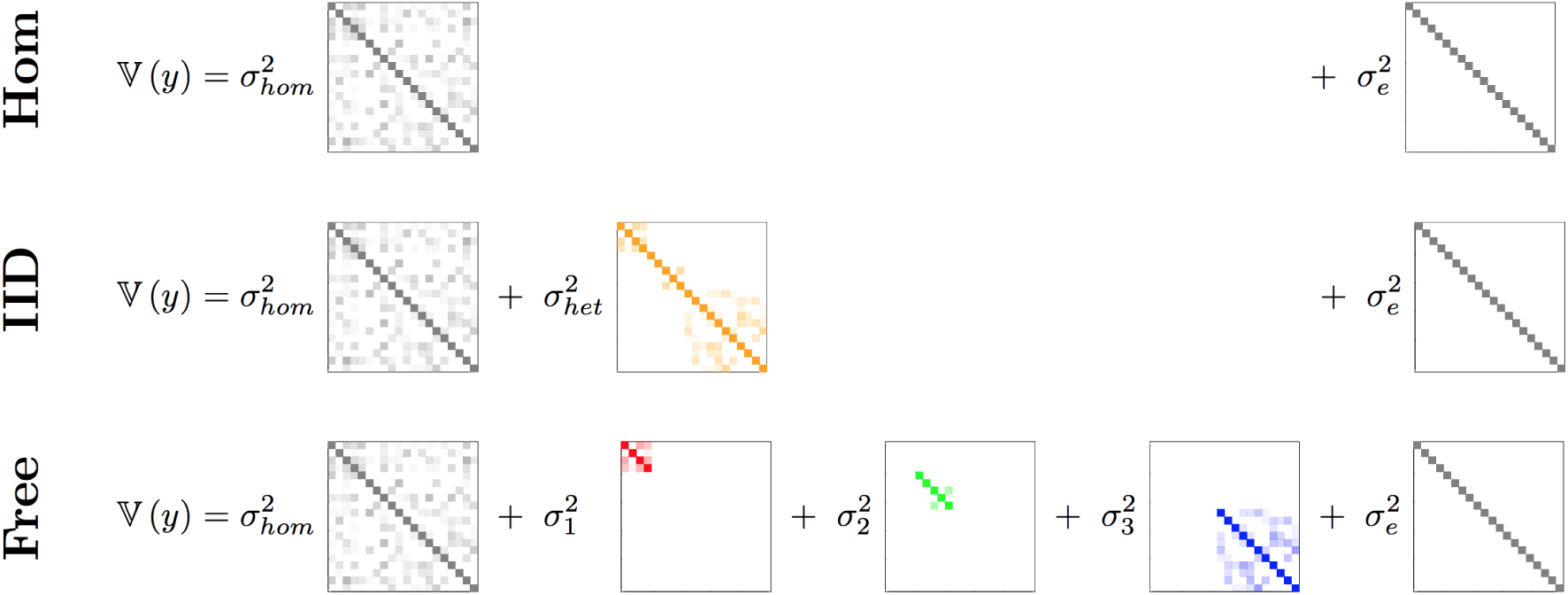
Depiction of the covariance matrices fit by the three LMMs we study. Samples are grouped by membership in one of three environments (red, blue and green). The ordinary GREML model (top) fits the additive heritability (governed by 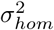, in grey) and noise 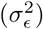. Darker colors indicate absolute values closer to 1, the value for all diagonal entries. The IID GxEMM (middle) adds a heterogeneous effect that modifies heritability only within environments (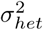, orange). The Free GxE mixed model (bottom) allows different per-environment heritabilities 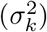.

We introduce ordinary genetic linear mixed models in Section 2.1. We then generalize them to model genetic interactions, without imposing the simplifying assumptions used in previous work, in Section 2.2. We discuss issues of restricting the parameter space, identifiability, scaling, testing, and defining environment-specific heritability in Sections 2.3-5. We then show GxEMM delivers roughly unbiased estimates and calibrated tests in simulations from the GxEMM model, and also that GxEMM can increase power to detect heterogeneity over existing tests. Finally, to demonstrate the practical utility of GxEMM, we re-analyze an MD dataset and validate our previous conjecture that major lifetime stress obscures the genetic impact on MD risk [16]. We conclude with comments on possible future methodological extensions and practical applications.

## 2 Polygenic GxE Mixed Model

### 2.1 Modelling polygenic additive effects

We assume a quantitative trait *y* ϵ ℝ^*N*^ measured on *N* samples. We allow *Q* background covariates in *X* ϵ ℝ^*N×Q*^. Our focus is on the *L* SNPs measured in the genotype matrix *G* ϵ ℝ^*N×L*^. We assume columns of *G* are scaled to mean zero, variance one.

With this standard notation, the additive linear mixed model can be written:

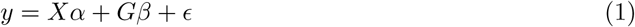

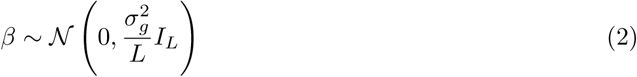

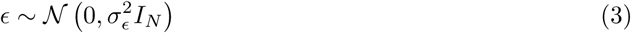

We assume *Q* ≪ *N* and spend the full *Q* degrees of freedom to estimate *α* as a fixed effect. In contrast, we model *β* as a random effect. This can be motivated as a genuine prior that all SNPs have small, nonzero effects, or as a convenient model that yields an accurate maximum likelihood estimate for ‖*β ‖* ^2^ under more general assumptions [32].

The normality assumption on *β* has the substantial practical advantage that ‖*β‖* can be estimated without calculating the high-dimensional *β*. This is accomplished by marginalizing out *β*, which gives a simpler and (essentially) equivalent formulation of the above mixed model:

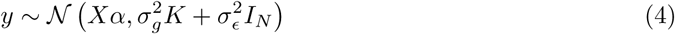

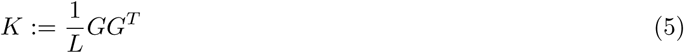

This defines *K* as the kinship/genetic relatedness matrix. Because *G* is assumed to have centered and scaled columns, *K* is a natural estimator of genetic similarity. While the model in (1) appears to require expensive computations in *β*, the marginalized model in (4) depends only on 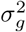 (and *α* and 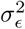). (4) can be fit by REML, which fits the variance components after projecting out *X*

One main use for LMMs in genetics is to estimate the additive, or homogeneous, or narrow-sense, heritability, which we define as 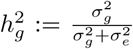 (assuming that *G* has centered and scaled columns). Conceptually, this quantifies the aggregate genetic contribution without identifying specific effects.

Another main use for LMMs in genetics is GWAS, accomplished by adding one SNP (at a time) as a column of *X* and testing if its corresponding entry in *α* is zero. The random genetic effect can be seen as adjusting for either polygenic background signal or non-genetic confounding [25]. A simple, common approximation to LMM-based GWAS is to only fit the variance components once, under the null hypothesis that SNPs have no effect [33], which works well in practice because SNP effects tend to be individually negligible (but see [34]).

### 2.2 Modelling polygenic interaction effects

We now assume *P* environmental variables have been measured in *Z* ϵ ℝ^*N×P*^. Like *X*, *Z* can contain binary and/or continuous values. We define the GxE mixed model (GxEMM) by adding environmental main effects and polygenic SNP-environment interactions to the ordinary LMM:

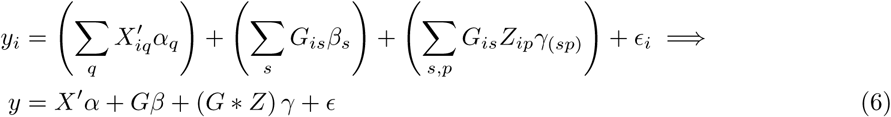

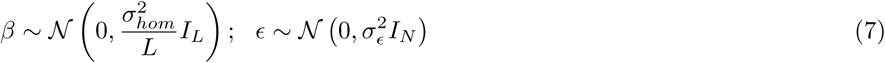

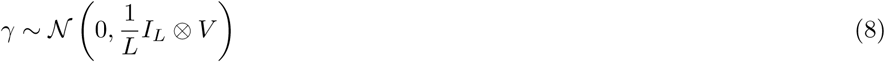

where ⊗ is the Kronecker product, * is the column-wise Khatri-Rao product, and *X′* are the fixed effects. We assume *P* ≪ *N* and estimate *Z* as a fixed effect, i.e. we define *X′* := (*X* : *Z*). We retain the spherical Gaussian priors on *β* and *ϵ* from the additive LMM, though we rewrite 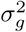 as 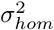 to emphasize that the interaction model permits heterogeneous sources of inheritance. We write *γ* _(*sp*)_ to indicate the entry of *γ* corresponding to the *s*-th SNP and *p*-th environment.

To our knowledge, the GxEMM model for the GxE coefficients, *γ*, is new. As for *β*, we assume that interaction effects are independent between SNPs, which is encoded in the assumed across-SNP covariance *I*_*L*_. However, we allow covariance between environments, in the same way for all SNPs, as encoded in the *P × P* across-environment covariance *V*. Unlike the *β* prior, which depends on one parameter, the *γ* prior depends on a matrix of parameters, *V*. Intuitively,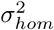 describes the size of *β*, while *V*_*pp*_ describes the size of the genetic interaction effects with environment *p*. The off-diagonal terms, *V*_*pp*_*′*, account for genetic effects that are shared between environments *p* and *p′*.

Just as the random *β* can be marginalized for additive LMMs, the random *β* and *γ* in GxEMM can be marginalized. This gives the random-effect formulation:

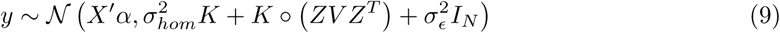

The GxE variance component decomposes into the Hadamard product (○) of genetic- and environmental-similarity matrices. This is actually if and only if: the only covariance matrices for *γ* that give (9) are of the form *I*_*L*_ ⊗*V* (assuming either large sample size or random *G*, Appendix).

The MLE for the model in (9), which involves a complicated covariance function of *V*, is not obvious. But, because the Hadamard product is linear, the matrix multiplications in *ZV Z*^*T*^ can be expanded and commuted through the Hadamard product with *K*, giving the equivalent expression:

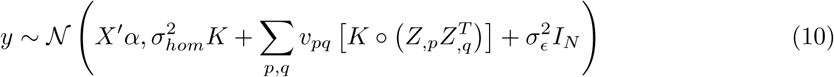

This covariance expression has a simple visual description for discrete environments (Figure 1).

(10) takes the form of a standard variance component estimation problem, which is useful because it can be solved by standard REML approaches. Specifically, we can use the GCTA software package [28] to fit Equation (10) with REML by using *X* as fixed effects and the similarity matrices 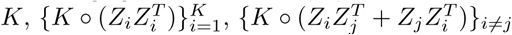, and *I*_*N*_.

Various restrictions reduce GxEMM to existing models. First, StructLMM is obtained if *L* = 1 SNP is used and 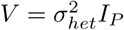, i.e. if interaction effects are i.i.d. across environments [31]. If, instead, *Z* contains only zeros and ones and 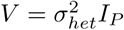, GxEMM becomes equivalent to the original GxE model in GCTA [28]. A related model was developed specifically for sex as the environment, which fits the sexes separately to allow sex-dependent genetic architecture, noise, and covariate effects [23]. GxEMM is most similar to iSet [29], except iSet focuses on low-rank *K*, assumes two discrete environments, and assumes samples are observed in all environments.

### 2.3 Restricting *V*

We never fit GxEMM with a general *V* in this paper because it has *O*(*P*^2^) parameters, which is computationally intensive for generic REML algorithms. We instead consider two nested restrictions to the model. First, Free GxEMM restricts *V* to be diagonal, i.e. 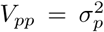 and *V*_*pq*_ = 0 for *p* ≠ *q*. While this imposes independence between environment-specific effects–after accounting for homogeneous effects–it allows the GxE heritability explained to vary freely between factors in *Z*.

Second, we evaluate an even simpler IID GxEMM that restricts 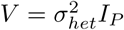, imposing that all columns of *Z* have similar genome-wide interaction effects. In particular, if *Z* contains unscaled discrete environment proportions, the IID model enforces constant heritability across environments. If, further, *Z* contains only 0s and 1s, IID GxEMM reduces to the GxE model in [28].

Enforcing diagonal *V* also makes 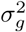 and *V* jointly identified. Otherwise, variance explained can be passed between 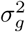 and *V* without changing the likelihood. To see this, let 1_*P*_ ϵ ℝ^*P*^ be a vector of 1s, let *≡* denote equal GxEMM likelihood, and assume, say, that *Z* has rows summing to 1. Then

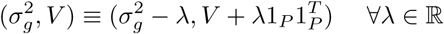

The constraint that *V* is diagonal, however, makes the right hand side infeasible for *λ* ≠ 0 (assuming *V* is feasible). Constraining 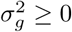 or *V* ⪰ 0, instead, would restrict *λ* to a proper interval of ℝ

When there are *P* discrete environments,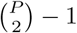 degrees of freedom are identified for *V*, while only *P -* 1 are identified for fixed effect interactions (per SNP). The extra degrees of freedom are identified by the prior on *γ*; e.g. when *V* is diagonal, the model prefers (*β, γ*) pairs where the average across-environment effects (per SNP) are explained by *β* rather than *γ*.

### 2.4 Centering and Scaling *G* and *Z*

*G* can be assumed mean zero WLOG because the full data likelihood is invariant under the mapping *G ↦ G* + 1_*N*_ *µ*^*T*^ for all *µ* ϵ ℝ^*L*^. *Z*, however, cannot be centered without changing the likelihood. This asymmetry between *Z* and *G* is because we treat *α* as fixed and *β* as random. The perturbation induced by centering *G* lives in the span of *Z*, and because *Z* is a fixed effect this perturbation is annihilated when fitting variance components by REML.

We assume columns of *G* have been normalized to empirical variance 1, corresponding to a particular assumption about the decay of effect sizes with MAF [35]. Almost nothing changes, however, if another (deterministic) column scaling is used, as long as 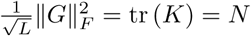

We do not assume *Z* is scaled to have columns with mean zero and variance one. Unlike with *G*, which has roughly exchangeable columns, columns of *Z* may have meaningful relative scales. We do, however, require that 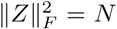, which comes without loss of generality and permits simpler formulas for heritability. For example, if environments are discrete and the data are restricted only to samples from environment *k*, the heritability in the new dataset would be

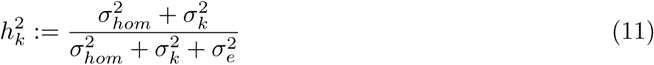

The full heritability in the observed data, accounting for both homogeneous and environment-specific heritability, replaces 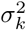 in the above equation with an average of the 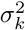 that is weighted by the size of each environment, 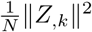.

The scale of columns of *Z* is irrelevant for Free GxEMM because the feasible set for *V* is closed under conjugation by diagonal matrices–multiplying *Z*_*,k*_ by *λ* can be counterbalanced by multiplying row *k* and column *k* of *V* by *λ*^-1^. However, the column scaling of *Z* affects results when *V* is forced to be spherical, as in IID GxEMM and existing GxE mixed models with *P >* 2.

### 2.5 Testing for significant GxE heritability

We use the standard LRT for GxE heritability terms and allow negative estimates for 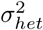 (and non-positive definite estimates for *V*) for three reasons. First, this reduces bias–provably so in the case of ordinary heritability [32]–which is important for aggregating estimates across traits. Second, if GxE heritability is defined as 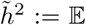, then the constraint 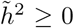 no longer requires either that 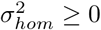 or that *V* ⪰ 0. Third, it is conceivable that environmental similarity could attenuate the importance of genetic similarity.

## 3 Simulations

Because genome-wide genetic data is complex, we validated GxEMM with simulations based on real genotypes from CONVERGE (see Section 4). For each simulation, we randomly choose *S* = 1, 000 SNPs to have causal additive and interaction effects and draw traits from the GxEMM model. We then fit three mixed models: the ordinary LMM with only a homogeneous genetic effect; the IID GxEMM with one parameter for GxE heritability; and the Free GxEMM allowing each heritability to vary freely between environments. Inside all mixed models, we use the standard genome-wide relatedness matrix as the causal SNPs are unknown in practice.

Specifically, we generate phenotypes from the following polygenic interaction model:

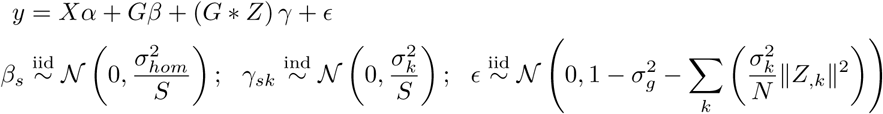

We define *X* to include the environments *Z* and the fixed effects we used in CONVERGE (Section 4) and draw 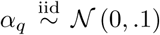 note that *X* is essentially irrelevant because it is projected out by REML. *G* contains the randomly chosen *S* causal SNPs. We scale columns of *X* and to *G* mean zero, variance one so that the effect sizes are easily interpretable. We choose 𝕍 (*ϵ*) so that the *σ*^2^ terms can be interpreted as fractions of (residual) phenotypic variance explained.

We define the environment *𝓏*_*i*_ for each person to be i.i.d. draws from one three discrete categories:

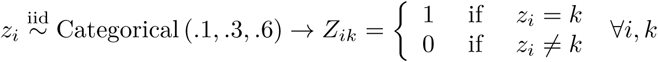

*→* means that we define *Z* by encoding the *K*-level factor *𝓏* as a binary matrix with *K* columns.

We varied the variance components in turn. First, we simulated under the homogeneous model by setting all 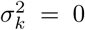 and varying 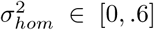 (Figure 2a,d). The IID and Free GxEMMs performed well in this null setting, each giving genetic heterogeneity estimates centered around zero and roughly calibrated tests (Figure 2a,d).

**Figure 2:**
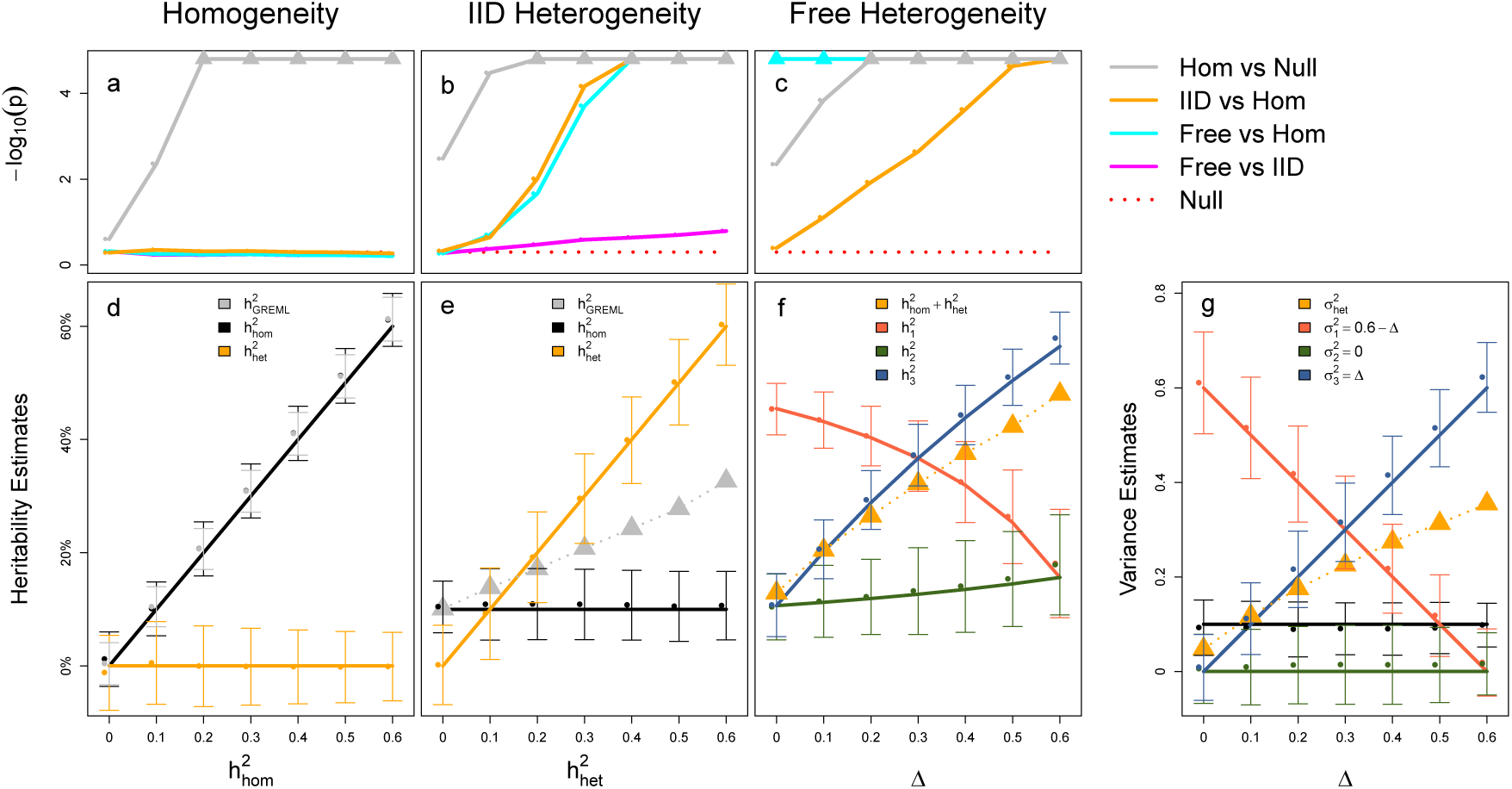
GxEMM tests (a-c) and genetic variance estimates (d-g) in polygenic GxE simulations with real genotypes. For each parameter setting, median -log_10_(*p*) across 100 independent simulations are shown for four LRTs under (a) genetic homogeneity, (b) constant genetic heterogeneity across environments, or (c) variable genetic heterogeneity. Large y-axis values are truncated as triangles (in c, pink and blue overlap). Null gives the median of 10,000 independent, null -log_10_(*p*) values. GxEMM estimates for the variance components are shown as medians (± one standard deviation). The ordinary heritabilities that would be obtained by restricting to one environment are shown in (g) by applying equation (11) to estimates from (f). For comparison, estimates from too-simple models are shown in triangles in (b,c).

To add IID genetic heterogeneity, we fix 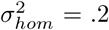 and vary 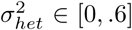 [0, .6] while requiring that 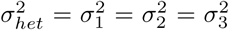 (Figure 2b,e). GxEMM accurately estimated the heterogeneous heritability, and the test for differences between the 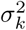 was roughly calibrated, especially for realistic values of genetic heterogeneity (‘Free vs IID’). Note that even though the Free model is needlessly general in this simulation, its test for genetic homogeneity only loses minimal power relative to IID GxEMM.

Finally, we fix 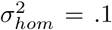 and add differences in the 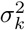 by varying a tradeoff parameter Δ ϵ [0, .6] (Figure 2c,f,g) We fix 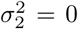 to reflect a moderately-sized subgroup without any specific genetic basis, and we interpolate the heterogeneous heritability between the small and large group by 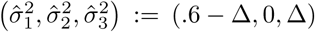. When Δ is small, most of heritability resides in the smallest environment, and the Free GxEMM test for genetic heterogeneity is substantially more powerful than existing, IID-based tests. The IID heterogeneity test gains power as Δ increases, meaning that we shift heritability from the small group to the large group, though Free GxEMM remains more powerful and explains more heritability. We also display the raw GxEMM estimates for 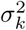 before rescaling to environment-specific heritabilities with equation (11) (Figure 2g).

## 4 Polygenic heterogeneity in major depression

Major depression is a moderately heritable trait (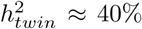 [36]) that is likely highly heterogeneous. The CONVERGE consortium [37] was established to enrich genetic signal for MD by recruiting only women with recurrent episodes of MD, between the ages of 30-60, and of Han Chinese ethnicity. Women were chosen because they have higher MD heritability (42%) than men (29%) [36]. Recurrent, clinically ascertained MD cases were selected because they likely harbor more heritable disease etiologies [38]. CONVERGE collected data from 5,303 cases and 5,337 matched controls. CONVERGE identified the first replicated GWAS results for MD, supporting the hypothesis that heterogeneity can mask causal genetic MD signals [37].

Further, in a direct test of this hypothesis, CONVERGE found three SNPs that are active only in MD cases without major stressful life events (SLE) [16]. We aim to use GxEMM to go beyond SNP-level tests by capturing differential genome-wide genetic architectures between SLE groups. This is motivated by analyses in [16] that found a suggestive and plausible but statistically insignificant difference in heritability between SLE groups.

We studied 9,303 samples with genotype and SLE data. We used an intercept, age, and SLE status as fixed effects. To control for population structure, we also included the top ten genetic PCs (from an LDAK relatedness matrix [39]) and their interactions with SLE [40].

We fit the homogeneous model and found 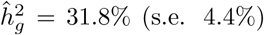. This estimate (and s.e.) is on the observed scale due to complexities of transforming to the liability scale in the presence of environmental main effects and differential prevalences. We next fit IID GxEMM and found 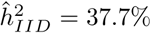 (s.e. 5.6%), which combines the homogeneous 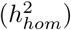 and IID heterogeneous 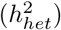 heritability components; we use the delta method to approximate the standard error. Finally, we fit Free GxEMM and found 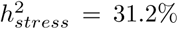 (s.e. 6.5%) and 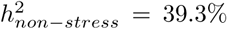 (s.e. 5.6%). The Free model fit better than the IID model (LRT *p*=0.00018), showing that the stressed samples harbor significantly lower genetic signal than the non-stressed.

[16] found three SNPs that interacted with SLE (rs7526682, rs11577545, and rs950893). To estimate the aggregate size of GxE effects that remain unaccounted for, we refit Free GxEMM while including these SNPs as fixed effects. While the stressed heritability estimate is basically unchanged (31.2%), the non-stressed heritability slightly decreases (38.0%). This trend suggests these three SNPs primarily act in non-stressed samples and that many GxE effects remain unexplained.

A significant caveat is that our current GxEMM implementation models binary trait heritability on the observed scale, treating the trait as quantitative rather than translating to the more natural liability scale. This translation is important even in the homogeneous case, as standard LMM gives biased heritability estimates for binary traits [41]. Further, even after translation, REML estimates are still biased, especially with preferential case recruitment and large covariate effects, and moment-based approaches are required instead [42, 43]. However, these methods are not easily applied under genetic heterogeneity, and REML is a common method in practice. We will extend GxEMM to more carefully handle binary traits and ascertainment in future work.

## 5 Discussion

Gene-environment interactions and small, genome-wide effects are separately well-documented. We unified approaches to study these two types of genetic effects with a Khatri-Rao linear mixed model for polygenic GxE called GxEMM. Marginalizing the GxE effects gives a simple, visually intuitive variance component model that can be fit with existing software. We showed the importance of the generality offered by GxEMM in simulations where the assumptions of existing methods fail. We showed the practical significance of GxEMM by analyzing a human major depression dataset and showing, for the first time, that major lifetime stress obscures the genetic impact on MD.

GxEMM can be applied to any genome-wide datasets with relevant environmental measurements. It is particularly useful and interpretable when partitioning heritability amongst discrete environmental factors. Though we have used the word environment, our approach can be used for anything that putatively interacts with the genome, including other genotypes [44] or study indicators in a random effect meta-analysis [45, 46]. The significant practical utility of GxEMM is that environmental variables need not be categorical and that different amounts of heritability can be attributed to different environments.

GxEMM could also be used for GWAS. Models similar to IID GxEMM have been shown to improve power and calibration [40], suggesting the Free GxEMM can further improve results when the IID assumption fails. In fact, GxEMM GWAS would cost no more than standard LMM-based GWAS when approximating variance components by their null estimates, which is common practice.

The main limitation of our GxEMM implementation relative to other GxE mixed models is computation time. In special cases, the problem can be rewritten to roughly reduce the complexity to *O*(*N*) [29, 31], which can be a major advantage over our *O*(*N*^3^) implementation. Fundamentally, we do not exploit the structure of the GxE kinship matrices, which almost certainly enables faster computation. Methodologically, it would be interesting to model environment-specific variances, like iSet [29]. Noise heterogeneity can bias our current GxEMM implementation, though we could in principle add *K -* 1 relatedness matrices for environment-specific noise.

## Appendix: Marginalizing random interaction coefficients

The point of the proposition is that the set of Gaussian GxE coefficients with covariance *D* ⊗ describes essentially the same distributions as the set Hadamard kinship matrices (*GDG* ^*T*^) *˚* (*ZV Z* ^*T*^). The former are the natural model for polygenic GxE; the latter are easily fit by REML and have already been studied in special cases.

**Proposition 1.** *Assume that γ ∼ 𝒩* (0, Σ) ϵ ℝ ^*P×L*^*, that G* ϵ ℝ^*N×L*^ *has continuous, random entries, and that Z* ϵ ℝ^*N×P*^ *has rank P, with P < N. Write K* := *GDG*^*T*^ *for some fixed D. Then*

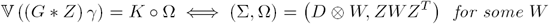

*where * is the column-wise Khatri-Rao product; ˚ is Hadamard; and* ⊗ *is Kronecker. Proof.* First, note the identity:

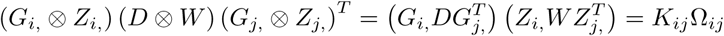

Now, right to left is easy:

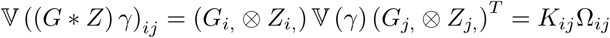

The other direction assumes decomposition of the variance into Hadamard products holds, so

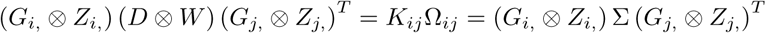

Using the standard identity (*A* ⊗ *B*) vec (*C*) = vec (*B*^*T*^ *CA*), where vec (·) concatenates columns of a matrix, the above equality can be written as

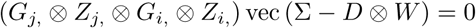

By assumption, this identity holds ∀*i, j* and almost all genotypes *G*_*i,*_*, G*_*j,*_ ϵ *R*^*L*^. The span of such pure four-way tensors is 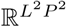, so its kernel has dimension zero and thus Σ - *D* ⊗ *W* = 0.

A similar result can be obtained if *N* grows large with fixed *L*.

